# COMPUTATIONAL COMPLEX PREDATOR-PREY DYNAMICS

**DOI:** 10.1101/151605

**Authors:** Adwitiya Chaudhuri, SK. Sarif Hassan

## Abstract

Two species predator-prey with stage structure of mature and immature mathematical models are studied over the last few decades. *Xin-an Zhang et al* studied a mathematical model with stage structure of two species in 2010. In this article, an attempt has been made to comprehend the coupled predator-prey dynamics with mature-immature stage structure and compare the dynamics with the existing model. The present model studied by *Xin-an Zhang et al* is purely realistic with assumptions of positive parameters. From the mathematical curiosity, we wonder to investigate the same with complex parameters and compared with the foreseen results. In addition, the present model is slightly modified to see some new dynamics of some additional fixed point including the previous fixed points.

## 1. Introduction

In mathematical ecology, the coexistence of species has become one of the interesting subjects of study [1, 2, 3, 4, 5, 6]. In the past few decades, dynamics of predator-prey have received great attention and have been investigated in a number of notable works [7, 8, 9, 10, 11, 12, 13]. There have been many modification of models from different perspectives such as introducing harvesting policy, control, delay and so on. The co-existence and stability of ecological systems has become one of the most prevalent phenomena and thus it is important for us to comprehend the permanence and global attractivity of systems [14, 15]. The strong persistence (permanent) and extinction are important concepts in predator-prey dynamics. In real systems, almost all species have their stage structure of mature and immature [16, 17]. In mature and immature species might also have different kinds/types such as active/inactive, diseased/nondiseased etc.

In the article [18], by *Xin-an Zhang et al*, the dynamics of the two stage structure of immature and mature predator-prey model is studied with the three assumptions as stated.

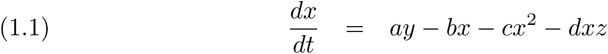

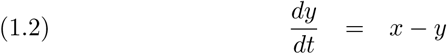

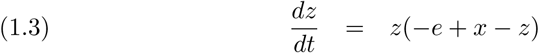

The parameters *a, b, c, d* and *e* are all positive constants. The variables in the system of eqs. (1.1 − 1.3) are defined as follows:

- *x*(*t*) denotes the population size of immature prey at time *t.*
- *y*(*t*) denotes the population size of mature prey at time *t.*
- *z*(*t*) denotes the population size of predator at time *t.*

The system of eqs. (1.1 − 1.3) is transformed from the original model

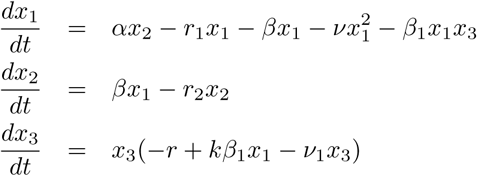

where *α*, *r*_1_*, r*_2_, *β*, *β*_1_, *v, v*_1_, *r* and *k* are all positive constants, *k* is a digesting constant. The transformed parameters *a*, *b*, *c*, *d*, *e* are associated as 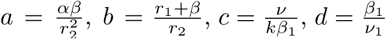 and 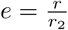. The assumptions to the original model as stated in the article [18] are given by

- *H*_1_: The birth rate of the immature population is simple-varied (proportional) to the existing mature population with a proportionality constant *α*; for the immature population, the death rate and transformation rate of mature are simple-varied to the existing immature population with proportionality constant *r*_1_ and *β*; the immature population is density restriction 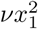.
- *H*_2_: The death rate of the mature population is simple-varied to the existing mature population with a proportionality constant *r*_2_.
- *H*_3_: The second species is a predator of the immature population of the first species; the second species satisfies the logistic predator-prey model.

We respect all the assumptions as stated (*H*_1_, *H*_2_ and *H*_3_) with a jump to the notion of *coupled dynamics* of prey-predator which is defined as follows.

- By coupling of constituent variables *x*, *y* and *z*, we mean to consider them as complex variables (the real part and imaginary part) instead of just real. The complex variables *x*, *y* and *z* essentially refer simultaneous population of two different kinds (such as sexually active/inactive, diseased/nondiseased etc).
- The parameters involved in the system of nonlinear eqs. (1.1 − 1.3) are *a*, *b*, *c*, *d* and *e* also taken as complex numbers. The real and imaginary part of any of these parameters are associated to the two different kinds of prey (immature and mature) and predator. It is worth noting that consideration of complex parameters in the model would possibly loose biological direct implications but certainly this mathematical model would be mathematically complex and crazy enough which is one of the aims of the article to understand.

The main results of the article [18] are given below while the parameters and system taken real numbers as presented.

- The system of eqs. (1.1 – 1.3) has a positive equilibrium if and only if *a* > *b*+ *ce*
- If *a* > *b* + *ce,* then the positive equilibrium of the system is globally asymptotically stable.
- If *b* < *a* ≤ *b*+*ce*, then the non-negative equilibrium of the system is globally asymptotically stable.
- If *a* ≤ *b*, then the origin of the system is globally asymptotically stable.

Given this brief introduction, we proceed to the main content and before coming to the asymptotic behaviours of the prey and predator in the ecological system with coupled dynamical system, we find the discrete time system of the system of eqs.(1.1 − 1.3).

The discrete-time system of the system of eqs.(1.1 − 1.3) is given by

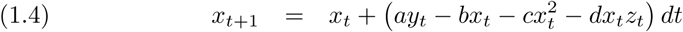

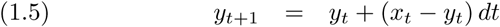

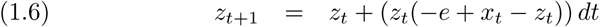

Here all parameters *a*, *b*, *c*, *d* and *e* are complex numbers and *dt* (which is considered to be 0.0005 throughout the rest of the article) is the delay term in discritizing the system.

The rest of the article is organized as follows. In *Sec. 2*, local asymptotic stability is discussed for all the fixed points of the system of *Eqs.*(1.4 − 1.6) and its modified system as discussed in the second last section.

## 2. Local Stability Analysis

Here we present two most well known necessary results in analyzing local asymptotic stability of a fixed point before we proceed to do so for the fixed points of the discrete time system of eqs.(1.1 − 1.3) [19].

### Result 2.1.

*Let* 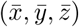 *be a fixed point of the system*

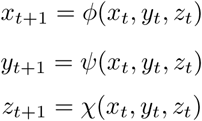

*Let* 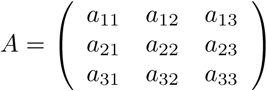 *be the jacobian at the point* 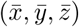 *with eigenvalues λ*_1_ *λ*_2_ *and λ*_3_*. Then:*

- 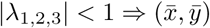 *is* locally asymptotically stable or attracting.
- *|Tr*(*A*) *|*< 1 + *Det*(*A*) < 2 *then* 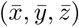 *is* locally asymptotically stable or attracting.
- *|λ_j_* | > 1 *for one* 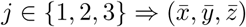 *is* repelling.
- *|λ_j_* | = 1 *for one* 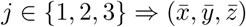 *is* saddle.

Another important result in understanding bound for zeroes of a polynomial with complex coefficients [20] is presented as follows.

### Result 2.2.

*Let p*(*z*) = *a*_3_*z*^3^ + *a*_2_*z*^2^ + *a*_1_*z* + *a*_0_ *be a cubic polynomial with complex coefficients such that for some h* > 0,

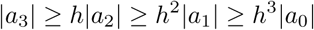

*Then p*(*z*) *has its all zeros in* 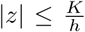 *where K is the greatest positive root of the biquadratic equation K*^4^ − *2K*^3^ + 1 = 0.

*Note that* 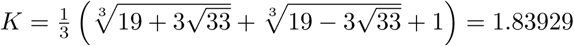

The fixed points 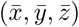 of the system Of eqs. (1.4 – 1.6) are solutions of the system of nonlinear equations:

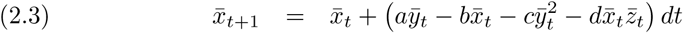

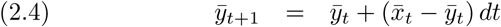

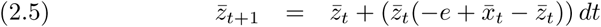

Consequently, the system of equations *Eqs.* (2.2 − 2.4) gives four unique fixed points

(0, 0, 0,) (0, 0, −*e*), 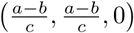 and 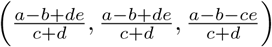. Before we proceed to study local asymptotical stability of the fixed points, we wish to raise few remarks about the fixed points as mentioned below:

- The fixed point (0, 0, 0) means that the predator-prey density will go on vanishing asymptotically. In an ecological system such a situation is not expected and it looks abolishing population and hence this fixed point is not of our interest.
- The other fixed point (0, 0, −*e*) signifies that the only second species (predator) will be permanent asymptotically and no mature and immature species (prey) will exist eventually which is also not an expected dynamics.
- The fixed point 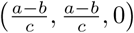 looks reasonable but again the second species (predator) is extinctive.
- The fixed point 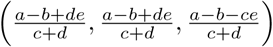 is an interesting from a healthy ecological population perspective since both the prey (immature and mature) and predator will exist permanently eventually as the system reaches equilibrium.
- It is to be noted that the eventually the density difference among immature, mature prey and predator is constant and that is *e.* For example, for the fixed point 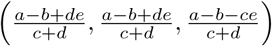 the asymptotic density of immature, mature prey and predator are 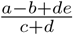,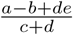 and 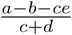 respectively. The density difference is 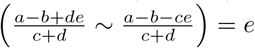 and the parameter *e* equals to 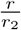. Among all the fixed points of the system, this is the only fixed point 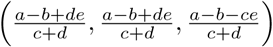 which is biologically significant and interesting since none of the three kinds of populations will get extinctive eventually as the system reaches its equilibrium.
- The system of *Eqs.*(1.4 – 1.6) does not possess any fixed points where only the immature prey will be permanent and mature prey and predator will be extinctive.
- The system of *Eqs.*(1.4 – 1.6) does not even have any fixed points where only the immature prey and predator will be permanent and mature prey will be extinctive.

We shall do now the local asymptotic stability analysis of these fixed points by making the system of *Eqs.*(1.4–1.6) linearized about the fixed points in the following subsections.

### 2.1. Local Stability Analysis of (0, 0, 0)

The linearized system *X_t_*_+1_ = *JX_t_* (where *X_t_* = [*x_t_*, *y_t_*, *z_t_*]*^T^* and *J* is the jacobian) is obtained by linearizing the model *Eqs.*(1.4 – 1.6) about the fixed point point (0, 0, 0). We wish to note that, we have considered *dt* = 0.0005 without loss of generality and proceed to apprehend the local stability of the fixed points.

Here the jacobian about the fixed point (0, 0, 0) while the discritizing delay term *dt* = 0.0005 is

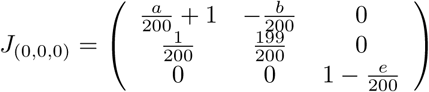

The characteristic equation of the jacobian matrix *J*_(0,0,0)_ is

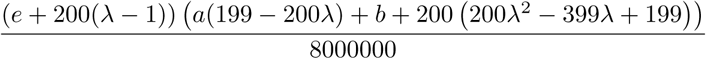

The fixed point (0, 0, 0) is attracting if all the zeroes lie inside the unit disk. Here we have the following theorem (follows from the *Result 2.2*) to ensure local asymptotic attracting behavior.

#### Theorem 2.6.

*The fixed point* (0, 0, 0) *is attracting if and only if there exists a constant h* > 1.83929 *such that*

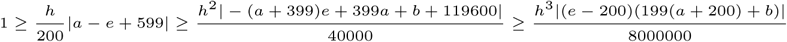

The eigenvalues of the jacobian matrix *J*_(0,0,0)_ are 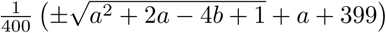 and 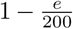. It is noted that, the *J*_(0,0,0)_ consists of only the parameters *a*, *b* and *e.*

Here we shall visualize three dimensional subspaces *S*_(*attracting*)_, *S*_(*repelling*)_ and *S*_(*saddle*)_ of 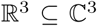 for the *dt* = 0.0005, which are shown in *Fig. 1.* In precise, *S*_(*attracting,dt*)_ denotes the space of parameters (*a*, *b*, *e*) in 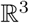 for which the fixed point (0, 0, 0) is *attracting* and similarly others.

**Figure 1.**
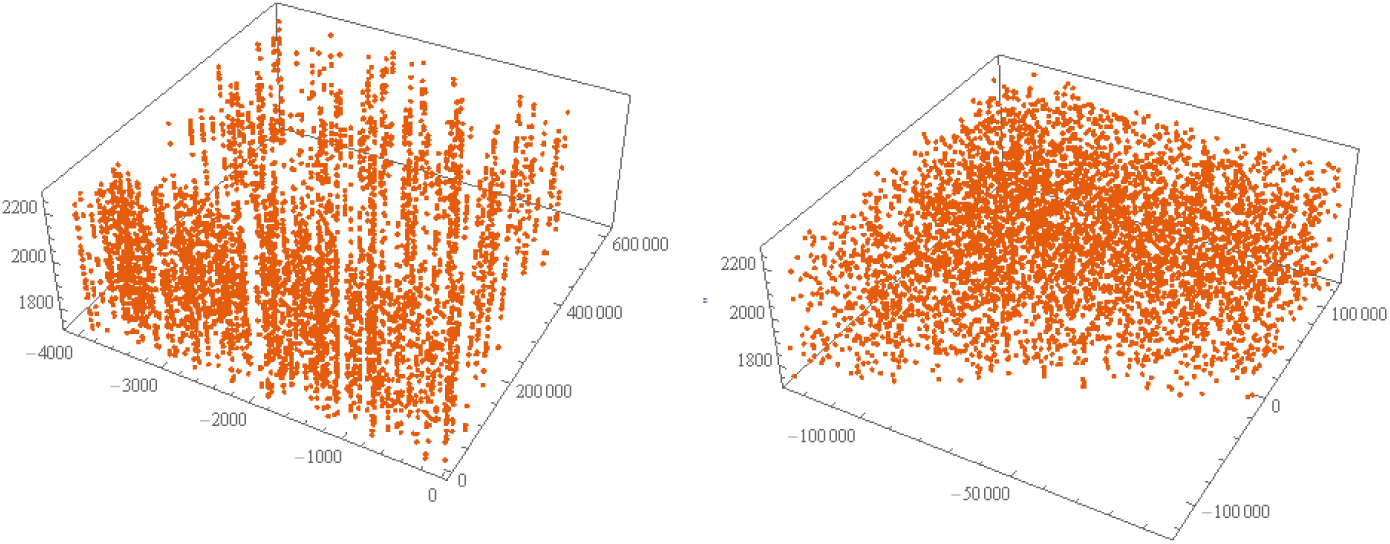
Left: *S*_(*attracting*),_ Middle: *S*_(*repelling*)_

It is to be noted that the set of parameters (*a*, *b*, *e*) such that the fixed point (0, 0, 0) is saddle is turned out to be *S*_(*saddle*)_ = {(−7999, 1.5992 × 10^7^, 1714.29), (−3999, −3999, 1714.29), (1, 1, 1714.29), (−7999, 1.5992 × 10^7^, 2285.71), (−3999., −3999., 2285.71), (1., 1., 2285.71)}.

We shall now see a set of examples of *attracting, repelling* and *saddle* behavior of the fixed point (0, 0, 0).

While *a* = 0.0326 + 0.5612*i*, *b* = 0.8819 + 0.6692*i*, *c* = 0.1904 + 0.3689*i*, *d* = 0.4607 + 0.9816*i* and *e* = 0.1564 + 0.8555*i*, then for ten different initial values taken from the neighbourhood of the fixed point (0, 0, 0), the trajectories are *attracting* to the fixed point (0, 0, 0) as shown in *Fig. 2.* The eigenvalues of the jacobian *J*_(0,0,0)_ are 0.996481 + 0.00571156*i*, 0.998682 − 0.00290556*i*, 0.999218 − 0.0042775*i* and all these eigenvalues lie inside the unit disk in the complex plane and hence the trajectories are attracting towards the fixed point (0, 0, 0). It is noted that the |*a*| ≤ |*b*|(0.562146 < 1.10706) is holding well, in the case of real positive parameters of the original model the condition foro global stability of the origin was *a* ≤ *b*.

**Figure 2.**
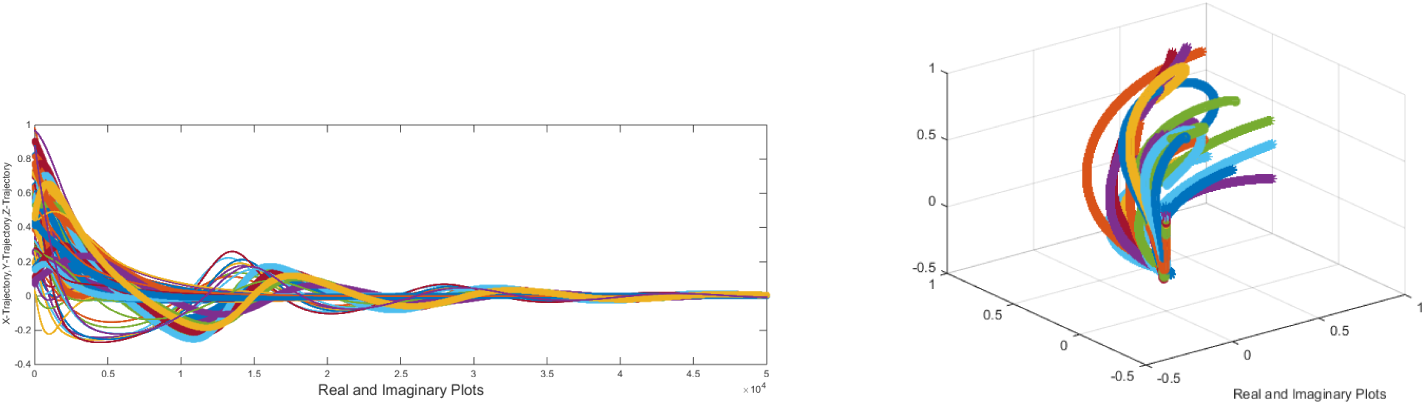
Attracting to (0, 0, 0) trajectories

While *a* = 0.8217 + 0.4299*i*, *b* = 0.8878 + 0.3912*i*, *c* = 0.7691 + 0.3968*i*, *d* = 0.8085 + 0.7551*i* and *e* = 0.3774 + 0.2160*i*, then for ten different initial values taken from the neighbourhood of the fixed point (0, 0, 0), the trajectories are *attracting* to the fixed point (0, 0, 0) as shown in *Fig. 3.* The eigenvalues of the jacobian *J*_(0,0,0)_ are 0.999551 − 0.00054046*i*, 0.999557 + 0.00268996*i*, 0.998113 − 0.00108*i* and all these eigenvalues lie inside the unit disk (modulus of each these eigenvalues are {0.999551, 0.999561, 0.998114}) in the complex plane and hence the trajectories are attracting towards the fixed point (0, 0, 0). It is noted that the |*a*| ≤ |*b*|(0.927364 < 0.970168) is holding here as well.

**Figure 3.**
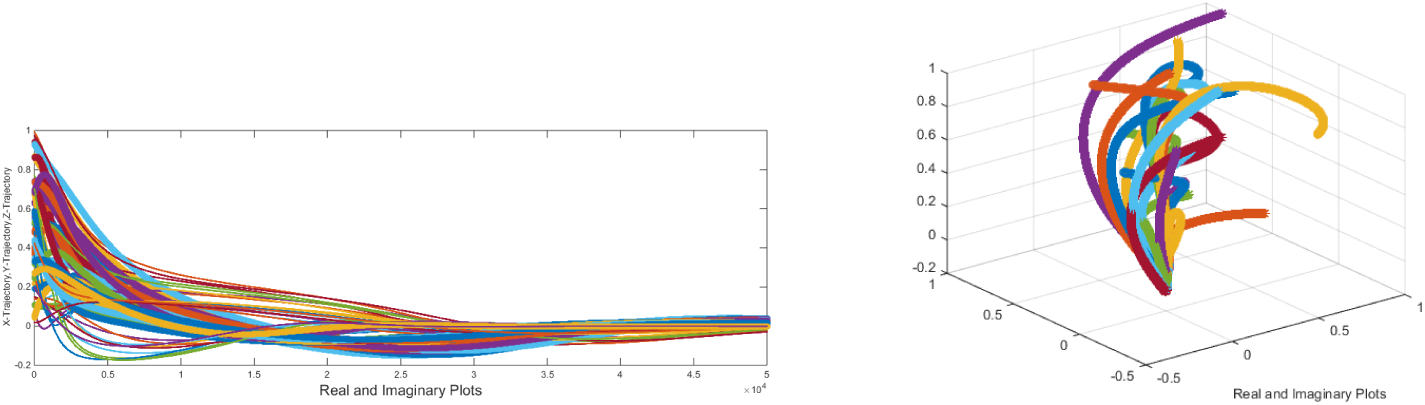
Attracting to (0, 0, 0) trajectories

#### Remark 2.7.

The fixed point (0, 0, 0) is *locally asymptotically stable* if |*a*| ≤ |*b*|.

In support of the *Remark 2.7*, we simulate a set of exemplary trajectories (twenty) for different set of parameters with |*a*| ≤ |*b*| for different initial values as shown in the *Fig. 4.*

**Figure 4.**
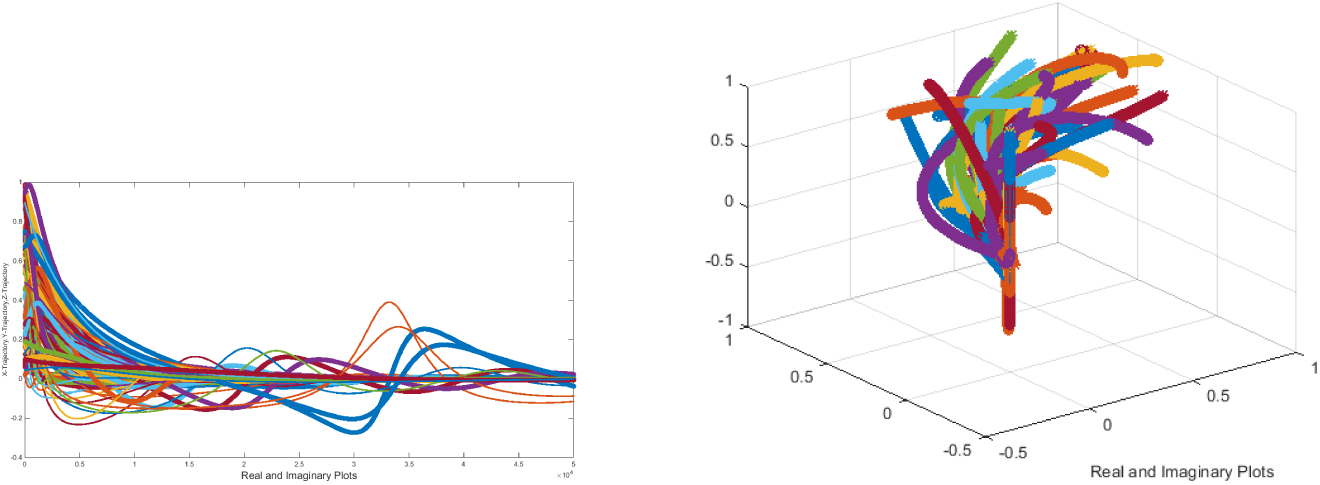
Attracting trajectories to (0, 0, 0) where |*a*| ≤ |*b*|

It is noted that, the converse of the *Remark 2.5* is not *necessary.* For counter examples, we have taken a set of twenty trajectories where the condition |*a*| ≤ |*b*| is violated (5*a* = *b*) and still trajectories are attracting towards the fixed point (0, 0, 0) as shown in *Fig. 5.*

**Figure 5.**
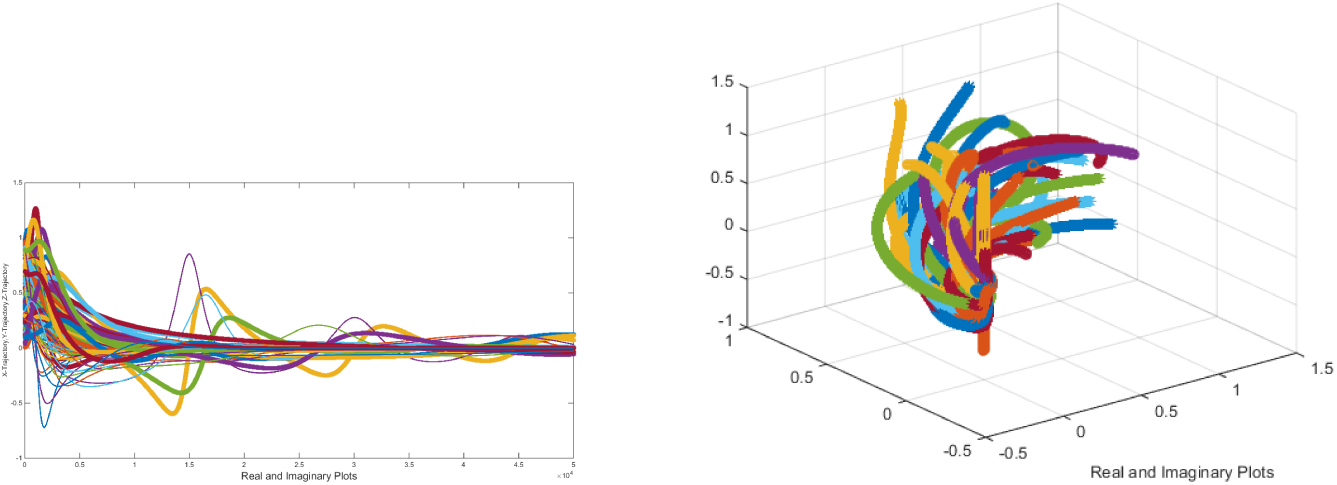
Attracting trajectories to (0, 0, 0) where |*a*| > |*b*|

Here we have taken an example of case where the fixed point (0, 0, 0) is saddle. Consider *a* = 1, *b* = 1, *c* = 0.6616 + 0.5170*i*, *d* = 0.1710 + 0.9386*i* and *e* = 2285.71 and then for any ten different set of initial values from the neighbourhood of (0, 0, 0), trajectories are away (converging to other fixed points) from the fixed point (0, 0, 0) (i.e. the fixed point is a saddle) as shown in the *Fig. 6.*

**Figure 6.**
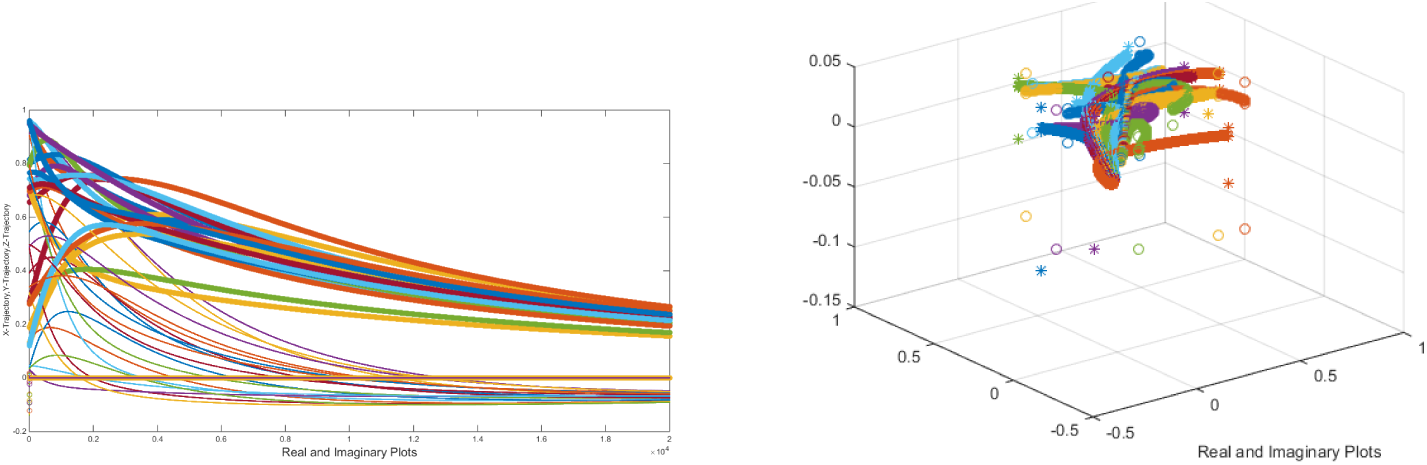
Saddle to (0, 0, 0) trajectories

Here we shall see few particular cases in this consequence. If the parameters *a* = 0 and *b* = 0, then the fixed point (0, 0, 0) cannot be attracted since one of the eigenvalues 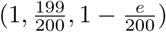 of the jacobian *J*_(0,0,0)_ is unity.

#### Remark 2.8.

While the parameter *b* = 0, then the fixed point is *attracting* if

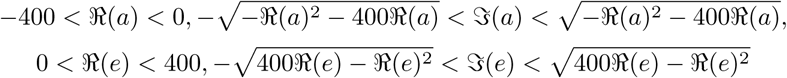

Note that, 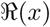 denotes the real part of the complex number *x.*

Such parameters *a* and *e* while *b* = 0 are plotted in the *Fig. 7.*

**Figure 7.**
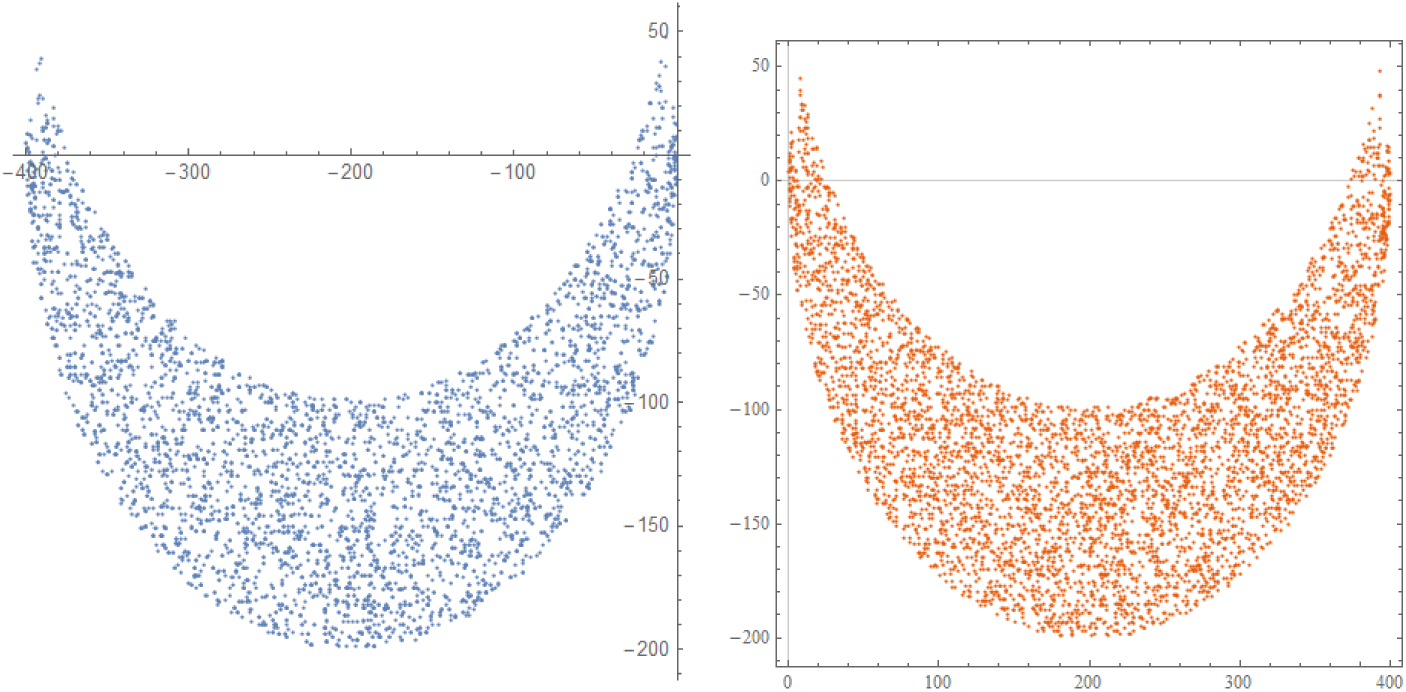
Parameter space of *a* (left) and *e* (right) for which the fixed point (0,0,0) is attracting

#### Remark 2.9.

If the parameter *b* = 0, then the fixed point is *repelling* if

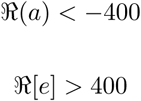

Such parameters *a* and *e* while *b* = 0 are figured out in the *Fig. 8.*

**Figure 8.**
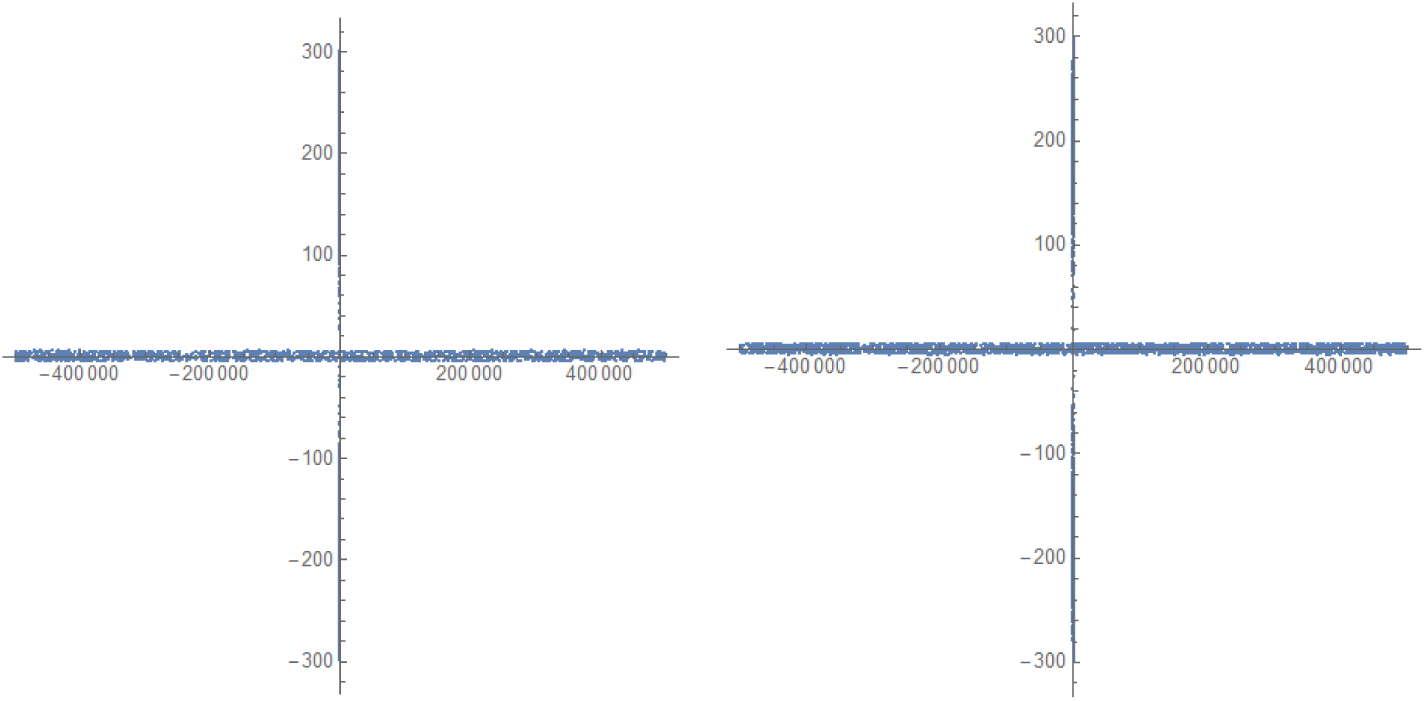
Parameter space of *a* (left) and *e* (right) for which the fixed point (0, 0, 0) is repelling.

### 2.2. Local Stability Analysis of (0, 0, −*e*)

The jacobian about the fixed point (0, 0, −*e*) while the discritizing delay *dt* = 0.0005 is given by

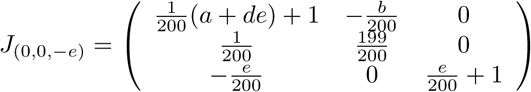

The fixed point (0, 0, −*e*) is attracting if all the zeroes lie inside the unit disk. The following theorem (follows from the *Result 2.2*) ensures the local asymptotic attracting behavior of the fixed point (0, 0, −*e*). The characteristic equation of the jacobian *J*_(0,0,−e)_ is

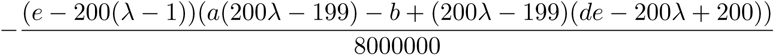

#### Theorem 2.10.

*The fixed point* (0, 0, −*e*) *is attracting if and only if there exists a constant h* > 1.83929 *such that*

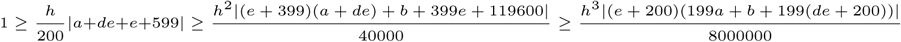

The eigenvalues of the jacobian *J*_(0,0,−*e*)_ are
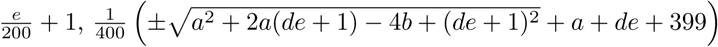.

#### Theorem 2.11.

*The fixed point* (0, 0, −*e*) *is repelling if*

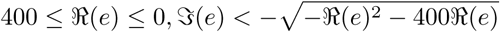

Here we are determine the sets of such parameter *e* such that the fixed point is *repelling and saddle* and such parameter are plotted in *Fig. 9* respectively. The parameter *e* lies in a circle as seen in the *Fig. 9.*

**Figure 9.**
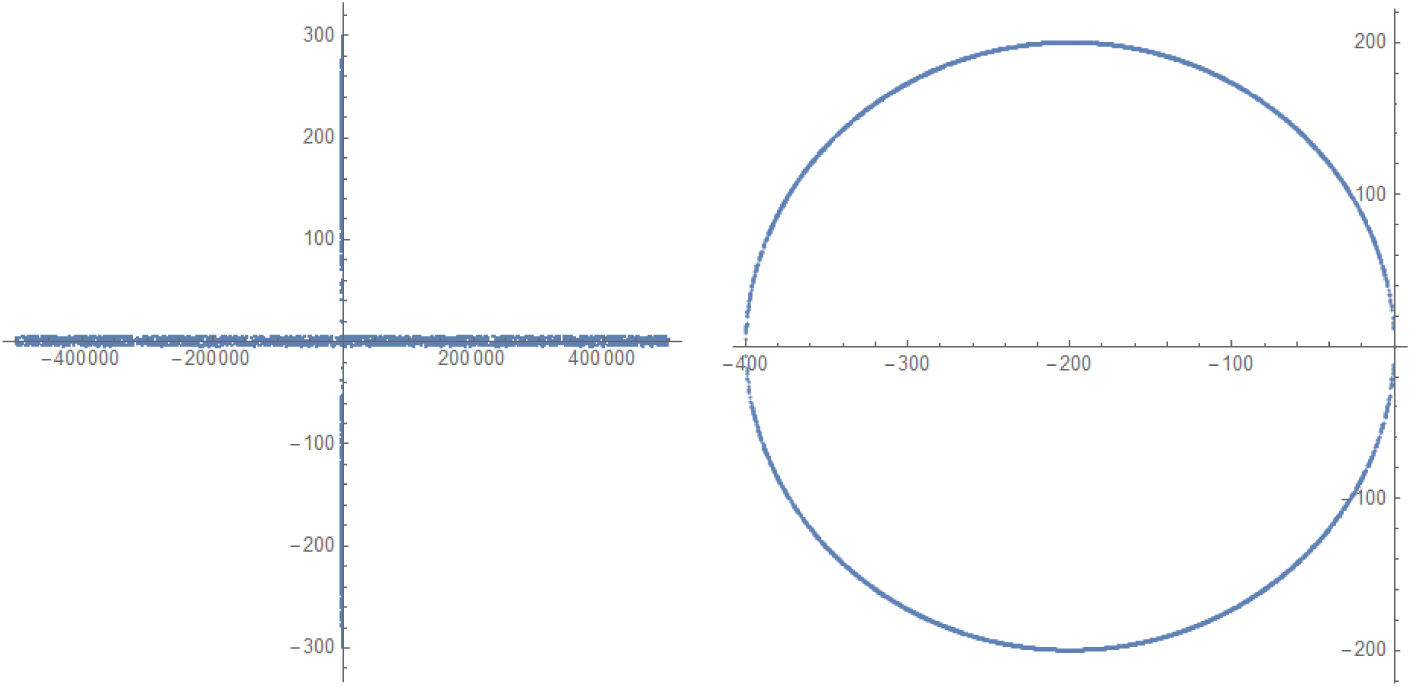
Parameter space of *e* (left: Repelling) and *e* (right: Saddle)

#### Theorem 2.12.

*The fixed point* (0, 0, −*e*) *is saddle if*

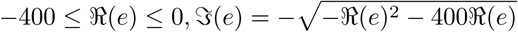

*Here* 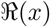 *and* 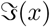 *are denoting real and imaginary part a complex number x.*

Here we have trajectories (repelling) from the fixed point (0, 0, −*e*) for different initial values and parameters as shown in *Fig. 10.* The trajectories are attracting to the other fixed point 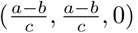.

**Figure 10.**
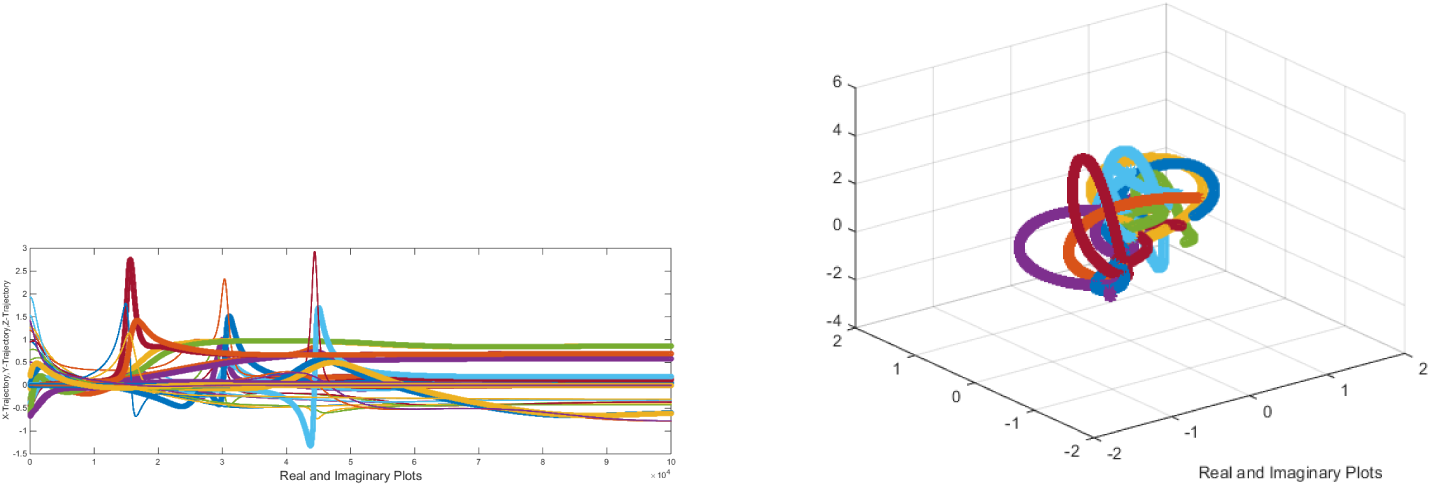
Repelling trajectories

It is worth noting that in an ecological system as mentioned in the beginning the fixed point (0, 0, −*e*) is not desirable. Computationally no complex parameters (except all are zero including h=0) is found for which the fixed point (0, 0, −*e*) is *attracting.*

### 2.3. Local Stability Analysis of 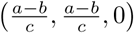

The jacobian about the fixed point 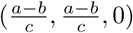 with *dt* = 0.0005 is given by

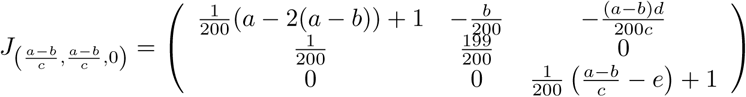

The characteristic polynomial of the jacobian 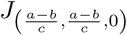 is

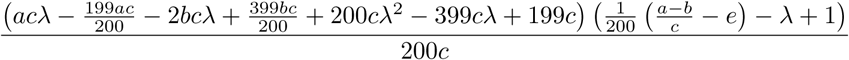

#### Theorem 2.13.

*The fixed point* 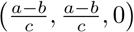 *is attracting if and only if there exists a constant h* > 1.83929 *such that*

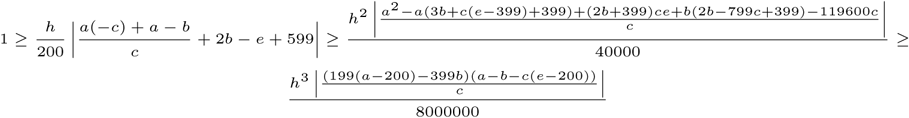

Here we illustrate an example of parameters such that the fixed point 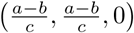 is attracting for different initial values. Consider 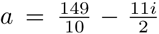, 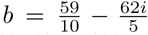, 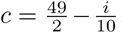, 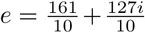, and then the fixed point becomes 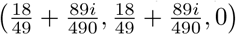 which is attracting for different initial values (20) taken from the neighbourhood of the point as shown in *Fig. 11.*

**Figure 11.**
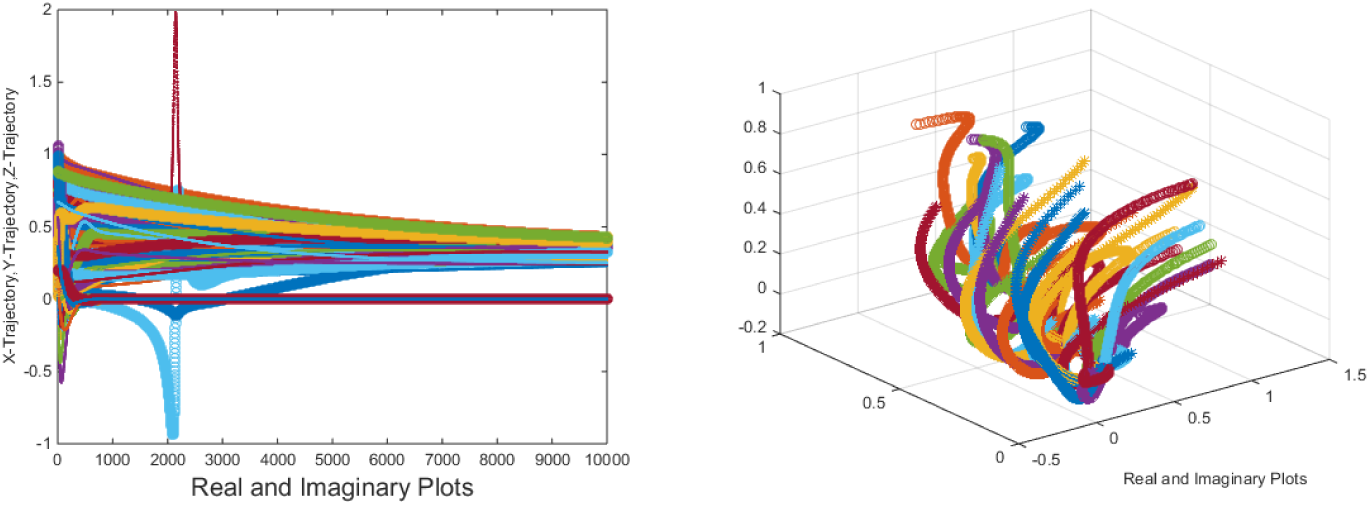
Trajectory of real and imaginary plots (left), 3D plot of the complex trajectories attracting to 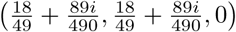

#### Theorem 2.14.

*If all the parameters a, b, c and e are real numbers and dt* = 0.0005, *then the fixed point* 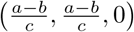 *is attracting if*

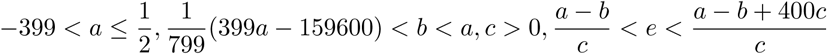

We have found a set of real parameters *a, b, c,* and *e* such that *Theorem 2.14* is valid. The parameters (*a*, *b*) and (*c*, *e*) are plotted in the figure *Fig. 12.*

**Figure 12.**
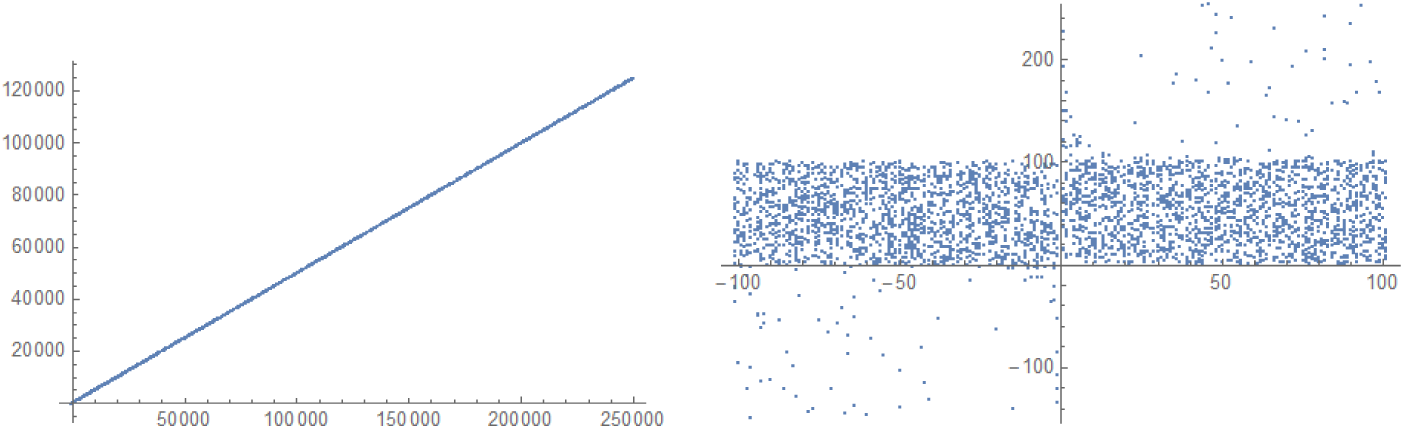
Two dimensional parameter spaces (*a*, *b*) (left) and (*c*, *e*) (right)

It is interesting to note that the parameters *a* and *b* are collinear and *c*, *e* are symmetric about the axes which is evident from the *Fig. 12.*

#### Theorem 2.15.

*If all the parameters a, b, c and e are real numbers and dt* = 0.0005, *then the fixed point* 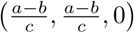 *is repelling if*

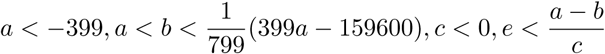

A set of real parameters *a, b, c,* and *e* is determined such that the fixed point 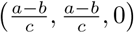 is *repelling.* The parameters (*a*, *b*) and (*c*, *e*) are plotted in the figure *Fig. 13.*

**Figure 13.**
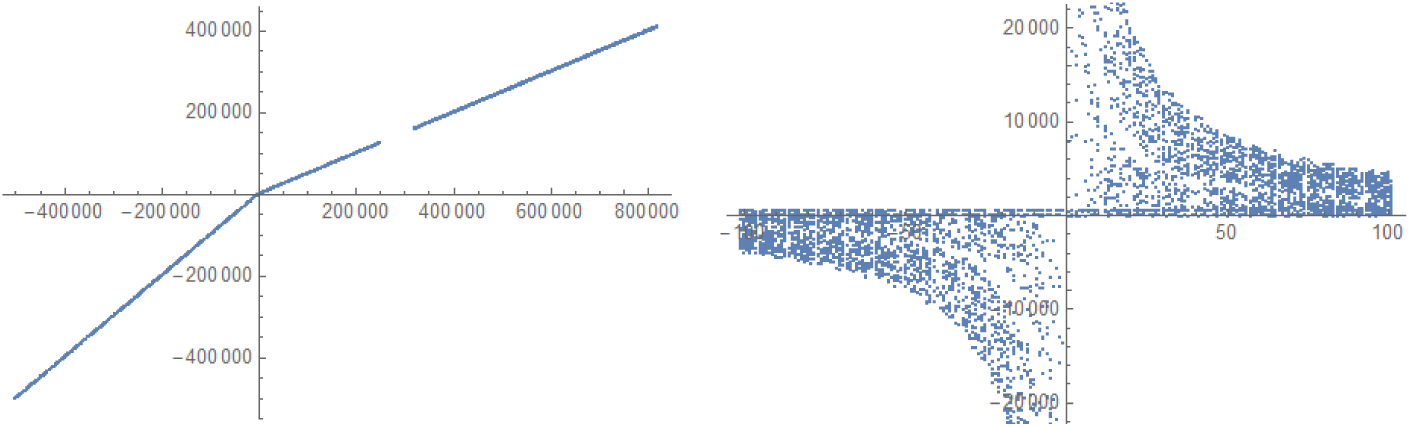
Two dimensional parameter spaces (*a, b*) (left) and (*c, e*) (right)

### 2.4. Local Stability Analysis of 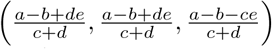

The jacobian about the fixed point 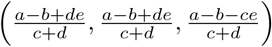 with *dt* = 0.0005 is given by

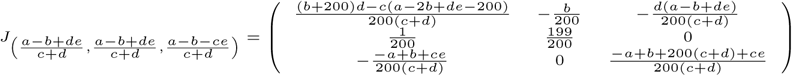

#### Theorem 2.16.

*The fixed point* 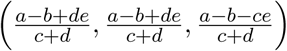 *is attracting if and only if there exists a constant h* > 1.83929 *such that*

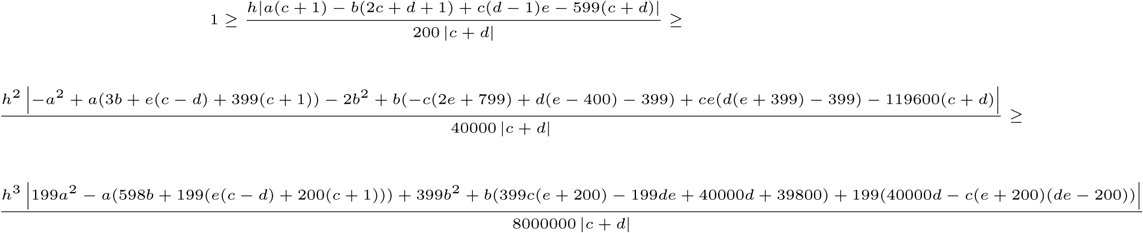

For arbitrary complex parameters *a*, *b*, *c*, *d* and *e*, it is difficult to apprehend the local behaviour of the fixed point and hence we see it through some particular cases.

First, we consider *b* = 0 and *e* = 0 and that essentially means that r2 ≠ 0, *r* = 0 and *β* + r1 = 0. That proportional constants (*r*_1_) and *β* of the death rate and transformation rate respectively of the mature prey to the existing immature prey are opposite in sign. Hence if the death rate of mature prey is positive then the transformation rate from immature prey to mature would be negative and vise versa. It may so happen that the parameters *r*_1_ and *β* both are zero and that is possible only if the transformation rate of mature do not proportional to the existing immature prey and such a situation comes when predation comes into action upon immature with a very high rate.

When the parameters *b* = 0 and *e* = 0 and then the eigenvalues of the jacobian 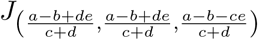 are 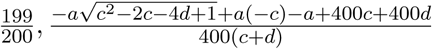, 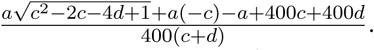.

In this consequences following results are obtained.

#### Theorem 2.17.

*For complex parameters with b* = *e* = 0 (*dt* = 0.0005), *the fixed point* 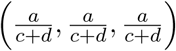 *is attracting if and only if there exists a constant h* > 1.83929 *such that*

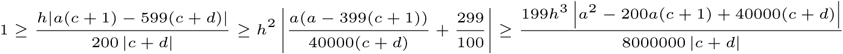

#### Theorem 2.18.

*For real parameters with b* = *e* = 0, *the fixed* 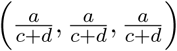 *point is attracting if*

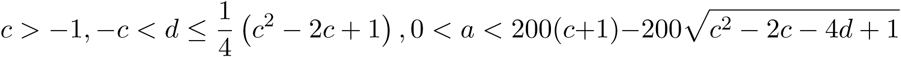

Here we present an example with different parameters (*b* = *e* = 0) such that the fixed point 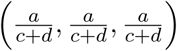 is attracting. The trajectory plots are given in the *Fig. 14.*

**Figure 14.**
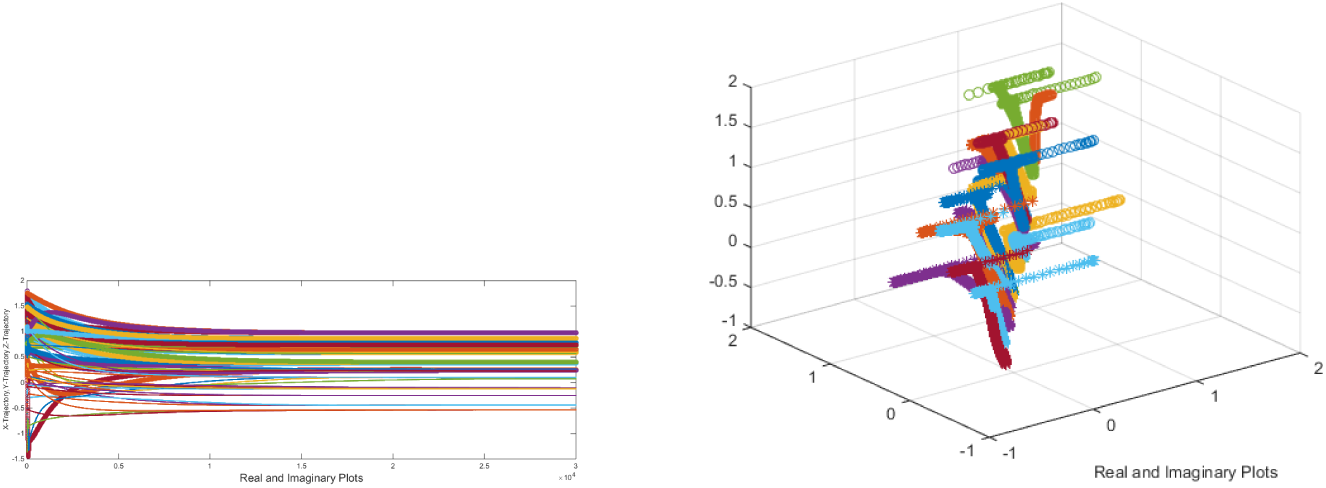
Trajectories converging to 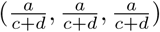

It is noted that the *Theorem-2.17* is shown to be valid for real parameters but it is seen in the example where complex parameters are taken which satisfy the condition

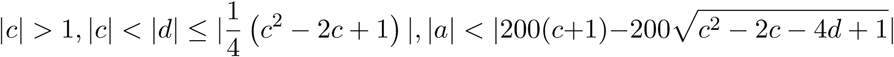

as stated in the *Fig. 14*, trajectories are attracting to 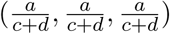 for ten different initial values as considered.

We have taken possible real parameters (*a*, *c*, *d*) such that the *Theorem 2.17* is holding well. The three dimensional plot of the parameters (*a*, *c*, *d*) is figured in *Fig. 15.*

**Figure 15.**
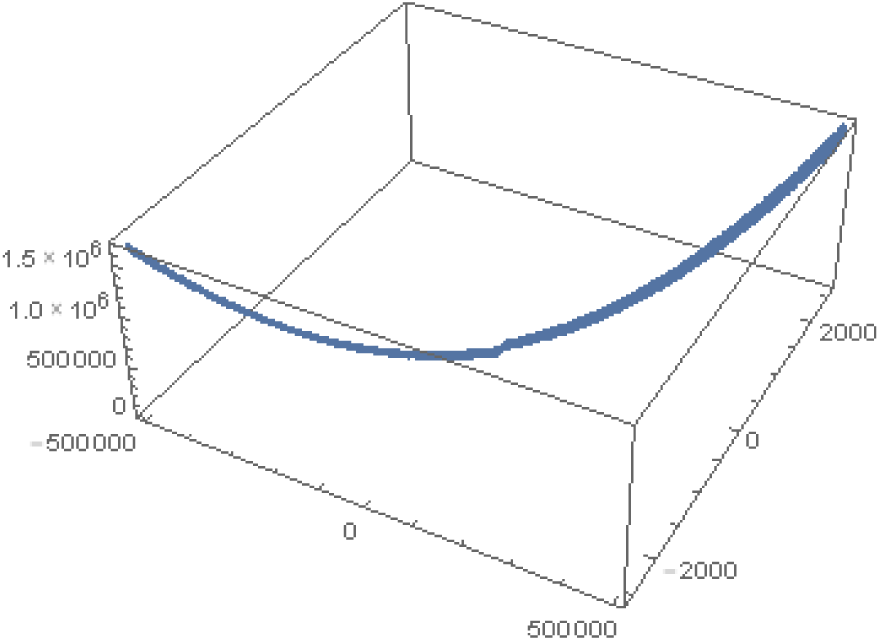
Three dimensional parameter spaces (*a, c, d*) such that 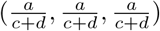 is attracting

It is noted that this is the only non-trivial desired fixed point where predator and prey are permanent with equal density in the ecological population.

#### Theorem 2.19.

*For real parameters with b* = *e* = 0, *the fixed point* 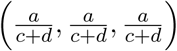 *is repelling if*

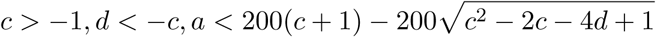

We have taken possible real parameters (*a*, *c*, *d*) such that the *Theorem 2.18* holds. The three dimensional plot of the parameters (*a*, *c*, *d*) is figured in *Fig. 16.*

**Figure 16.**
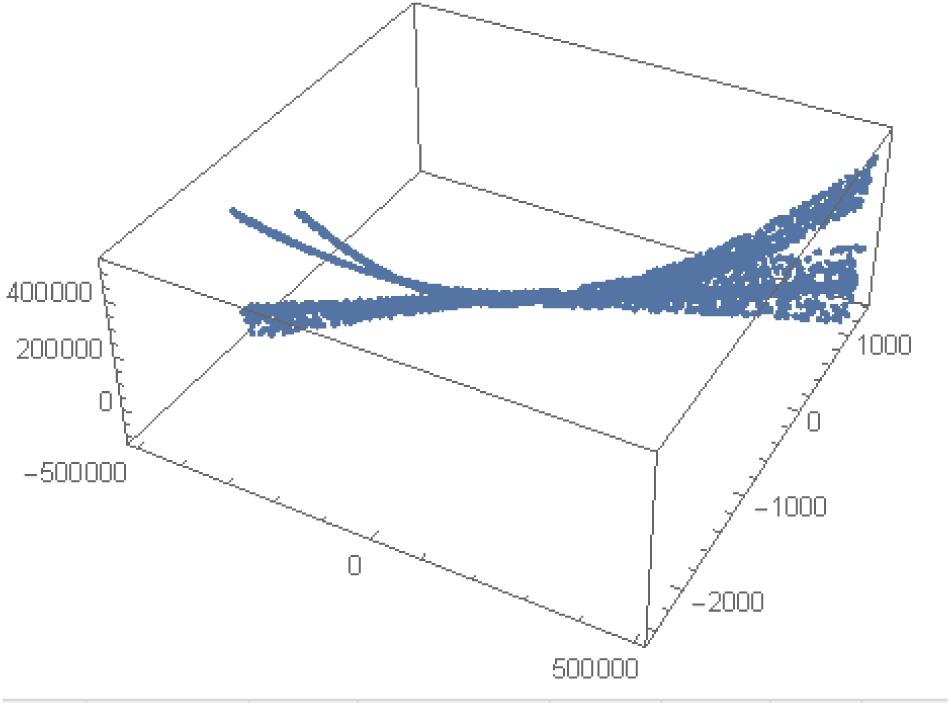
Three dimensional parameter spaces *(a, c, d)* such that 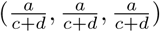 is repelling

#### Theorem 2.20.

*For real parameters a*, *c*, *d with b* = *e* = 0, *the fixed point* 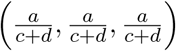 *is saddle if*

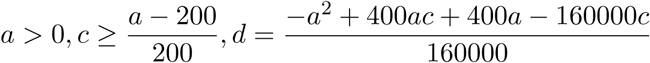

We have taken possible real parameters (*a*, *c*, *d*) such that the *Theorem 2.19* holds. The three dimensional plot of the parameters (*a*, *c*, *d*) is figured in *Fig. 17.*

**Figure 17.**
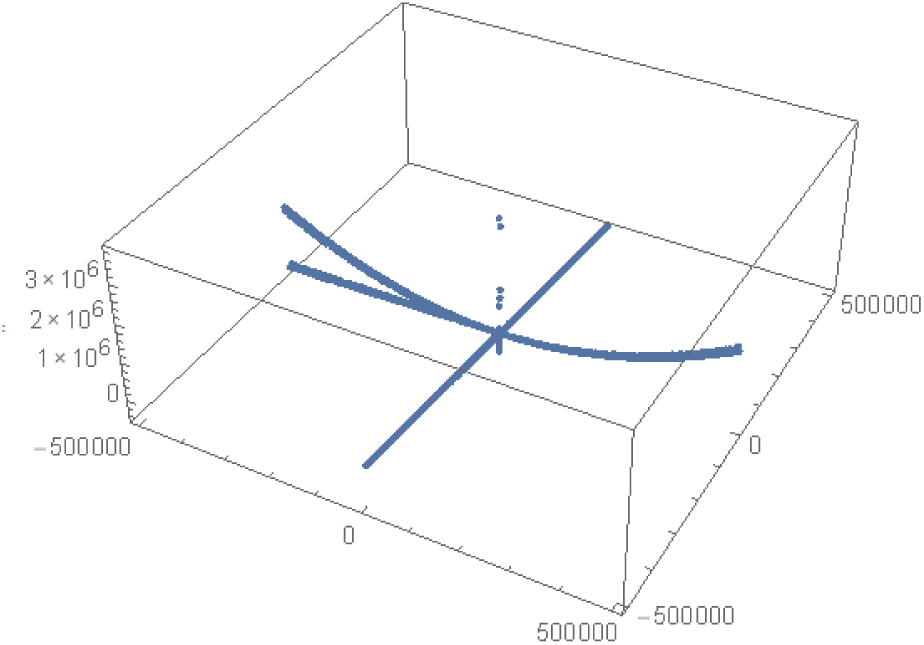
Three dimensional parameter spaces (*a*, *c*, *d*) such that 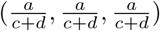 is saddle

If all the parameters *a*, *b*, *c*, *d* and *e* are equal then the parametric relation will be

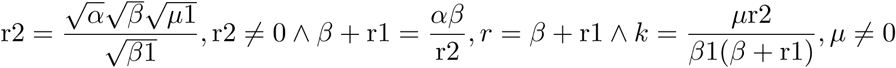

Here we assemble a set of parameters *α*, *β*, *β*_1_, *η*, *η*_1_, *k*, *r*,*r*_1_ and *r*_2_ which are implicitly associated to the parameters *a*, *b*, *c*, *d* and *e.* These complex parameters *α, β, β*_1_, *η, η*_1_, *k*, *r*, *r*_1_ and *r*_2_ are figured out in the following *Fig. 18.*

**Figure 18.**
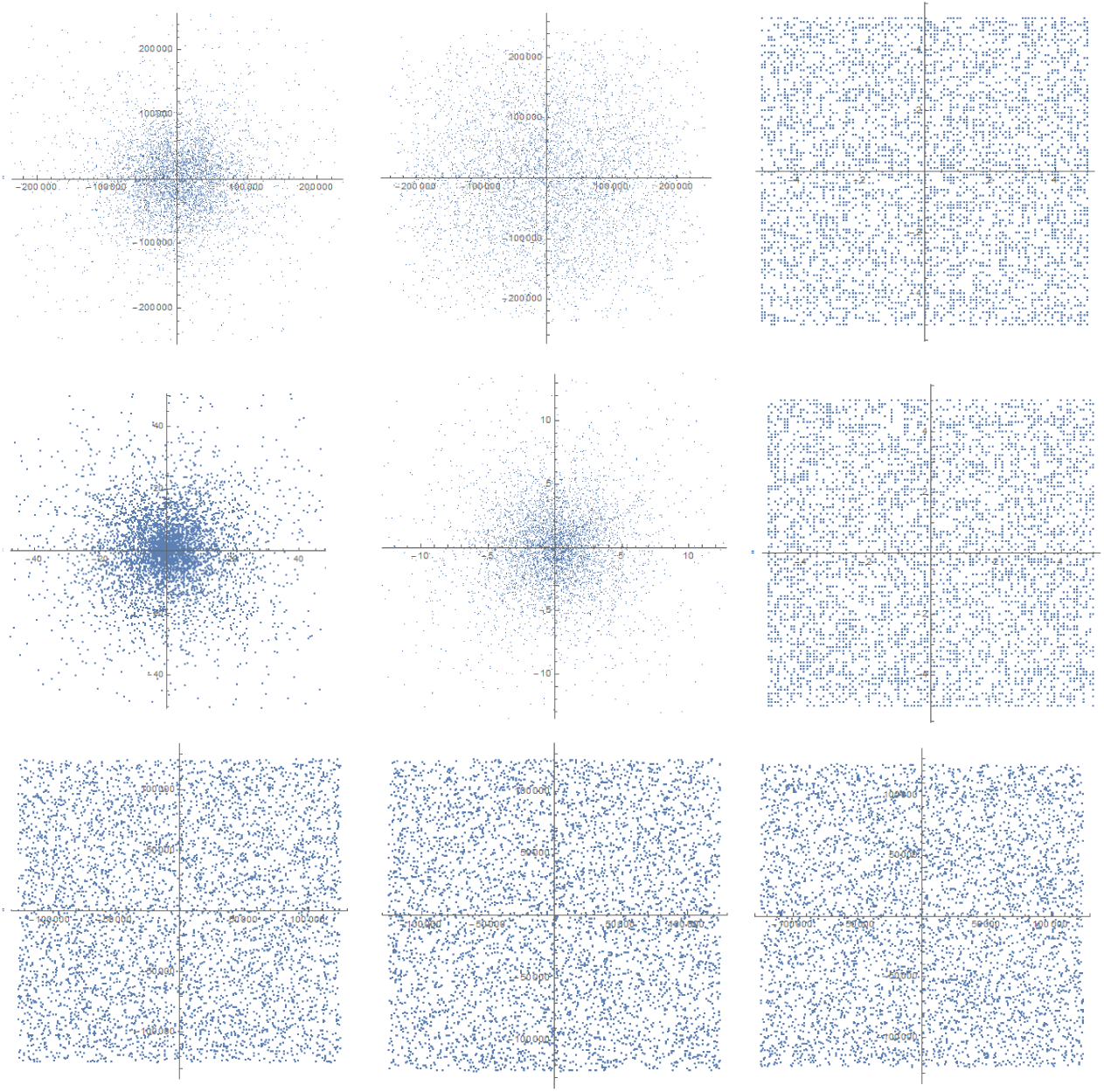
Complex parameters *α*, *β*, *β*_1_,*η*, *η*_1_*, k*, *r*, *r*_1_ and *r*_2_ such that the parameters *a*, *b*, *c*, *d* and *e* are all equal.

The existence of such parameters ensures the feasibility of the coupled preypredator local stability of special kinds which we shall explore immediately. When all the parameters *a*, *b*, *c*, *d* and *e* are same and equal to ħ then the fixed point becomes 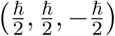 and let us explore local asymptotic stability.

#### Theorem 2.21.

*The fixed point* 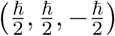 *is attracting if*

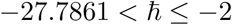

*or*

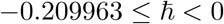

Here we illustrate the *Theorem-2.20* through an example which is shown in *Fig. 19.* We took *ħ* = −4 and for ten different initial values from the neighbourhood of the fixed point 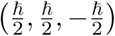, the trajectories are attracting to the fixed point (−2, −2, 2) as desired.

**Figure 19.**
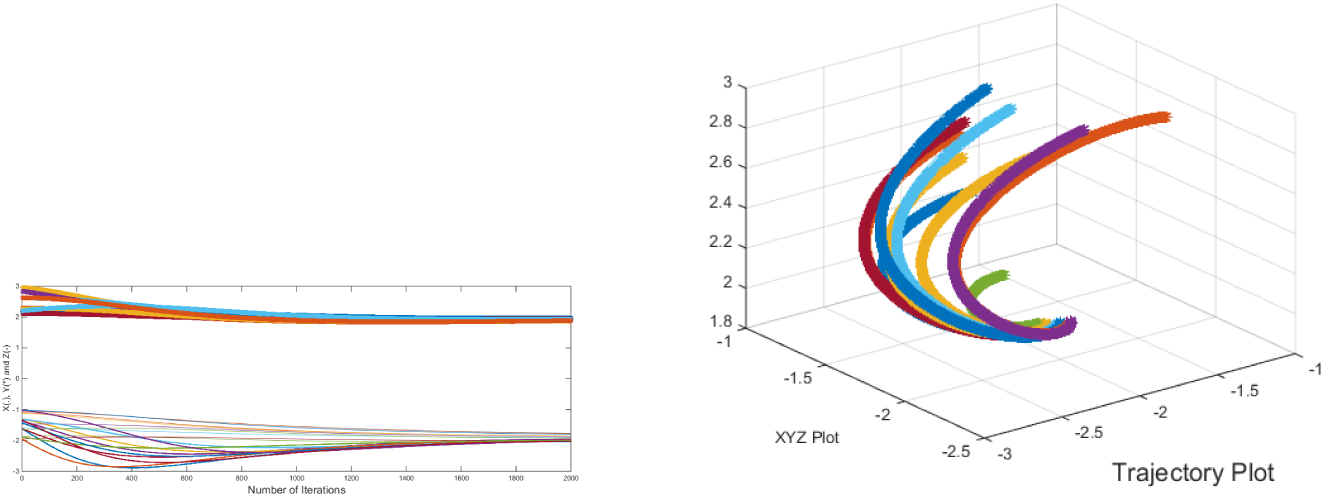
Trajectories attracting to (–2, –2, 2)

#### Theorem 2.22.

*The fixed point* 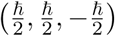 *is repelling if* 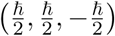 *if*

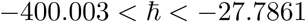

*or*

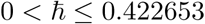

*or*

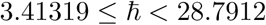

#### Theorem 2.23.

*The fixed point* 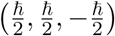 *is saddle if* 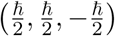 *if*

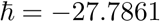

*or*

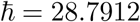

Here we illustrate the *Theorem-2.22* through an example which is shown in *Fig. 20.* We took *ħ* = 28.7912 and for ten different initial values from the neighbourhood of the fixed point 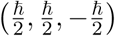, then the fixed point 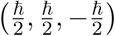 is a saddle as expected.

**Figure 20.**
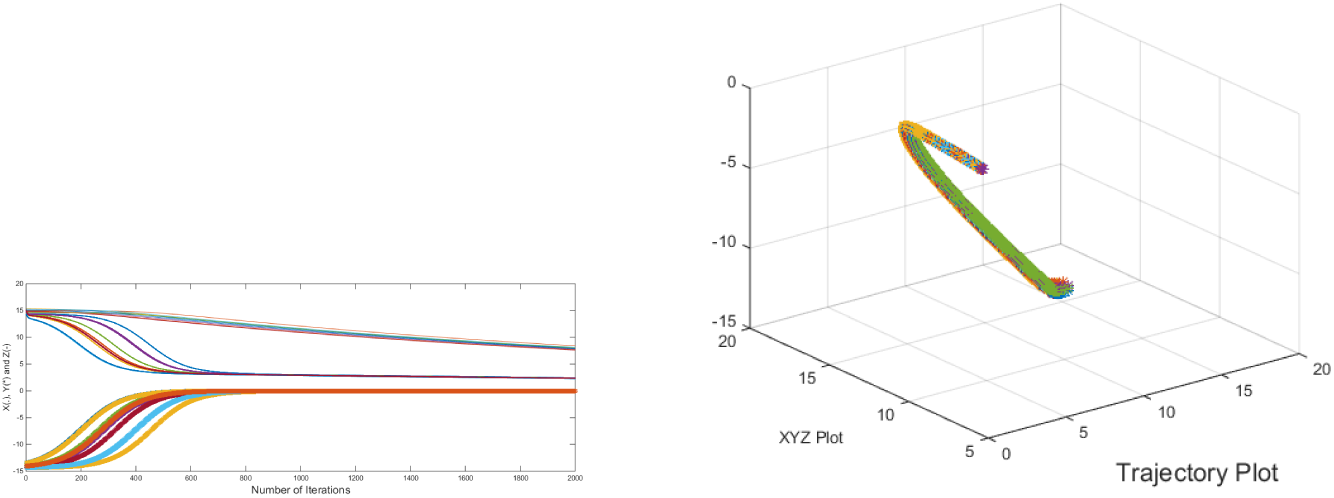
Trajectories attracting to (–2, –2, 2)

## 3. Modified System with Additional Dynamics

In this section, we modify the system of equations eqs.(1.1 – 1.3) such that the modified system will have all the previous fixed points including two additional fixed points of which, one kind is that only immature population will be permanent (rest are extinctive) and the other kind would be only immature prey and predator will be permanent ( only mature prey will be extinctive).

The discrete version of the modified model is given by …

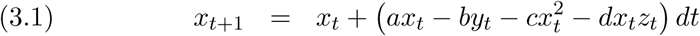

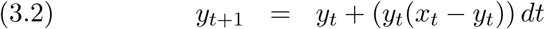

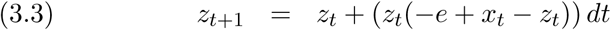

where all parameters *a*, *b*, *c*, *d* and *e* are complex numbers and *dt* is the delay term in discritizing the system. Note that here *a* and *b* parameters are swaped and accordingly the assumptions to the original model are swaped.

The fixed points of the system of eqs. (3.1 − 3.3) are (0, 0, 0), (0, 0, −*e*), 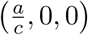,

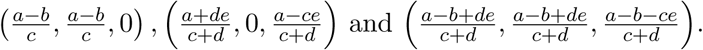

Including previous four fixed points of the system of eqs.(1.1-1.3), the other two additional fixed points are 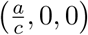 and 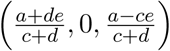.

Here the fixed point 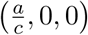 means that the immature species will be permanent and mature species including predator will be extinctive eventually.

The other fixed point 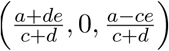 signifies that the only immature prey and predator will be permanent and mature species will be extinctive. These two eventual occurrence were missing in the system of eqs. (1.1–1.3) as mentioned in the remarks in the introduction section.

It is worth noting that the present system of eqs. (3.1 – 3.3) is enriched with additional dynamics including previous dynamics. Here we present local stability of these fixed points in the following subsections.

### 3.1. Local Asymptotic Stability of (0, 0, 0)

The jacobian about the fixed point (0, 0, 0) is 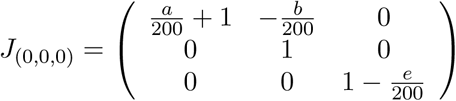, has three eigen-values which include 1 and hence the fixed point (0, 0, 0) cannot be *attracting.* The fixed point (0, 0, 0) is always a saddle as one of the eigenvalues is unity.

#### Theorem 3.4.

*The fixed point* (0, 0, 0) *of the system of eqs.* (3.1 – 3.3) *is repelling if*

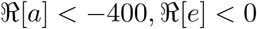

Here we present an example where the fixed point (0, 0, 0) is *repelling.* We consider the parameters *a* = 90 + 3.3*i*, *e* = 237.5 + 248*i* and *b*, *c*, *d* are arbitrary and then we found the trajectories for different initial values are *repelling* from (0, 0, 0) and *attracting* to the other fixed point(s) as shown in *Fig. 21.*

**Figure 21.**
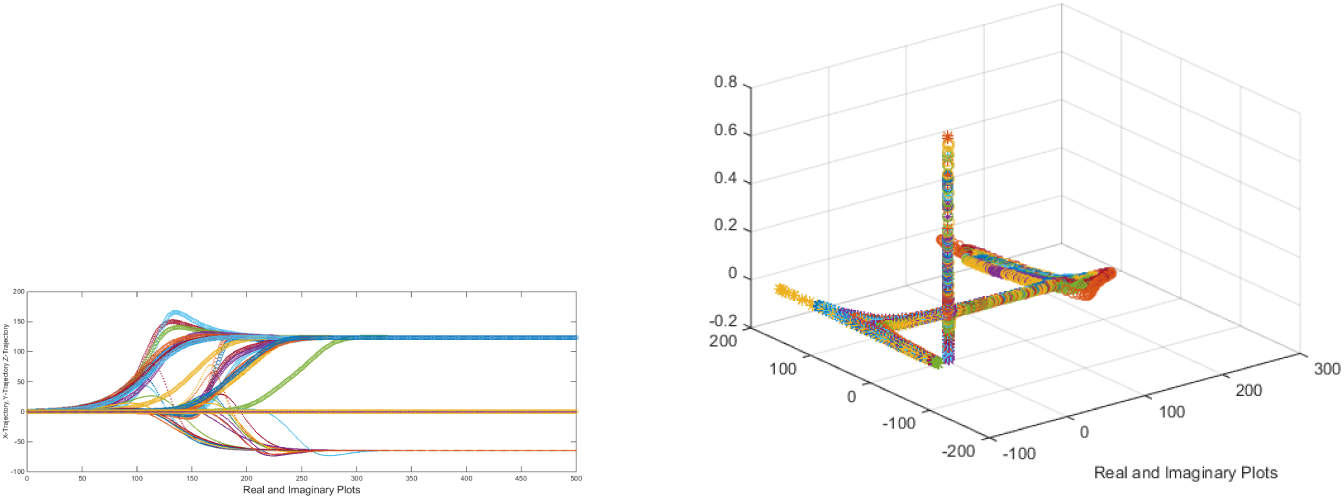
Trajectories repelling from (0, 0, 0)

### 3.2. Local Asymptotic Stability of (0, 0, −*e*)

One of the eigenvalues of the jacobian about the fixed point (0,0,-*e*) is unity and hence the fixed point would be either repeller or a saddle.

#### Theorem 3.5.

*For all the real parameters a, b, c, d and e, the fixed point* (0, 0, −*e*) *is repelling if*

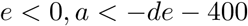

We found the space of real parameters (*a*, *d*, *e*) such that the fixed point (0, 0, −*e*) is repelling and the space is plotted in the *Fig. 22.*

**Figure 22.**
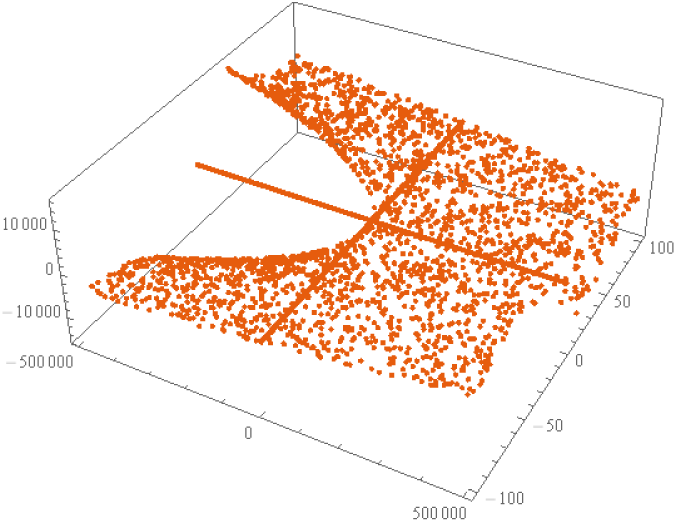
Trajectories repelling from (0, 0, −*e*)

### 3.3. Local Asymptotic Stability of the 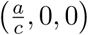

The jacobian about the fixed point 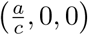 is

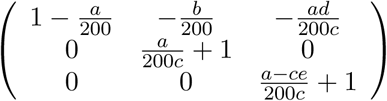

#### Theorem 3.6.

*The fixed point* 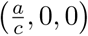 *is attracting if and only if there exists a constant h* > 1.83929 *such that*

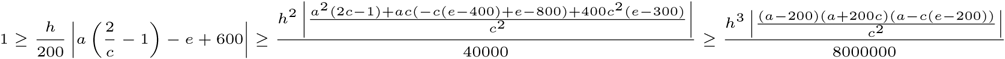

#### Theorem 3.7.

*If the parameters are real then the fixed point* 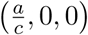 *is attracting if*

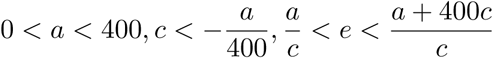

We found the parameters (*a*, *c*, *e*) with other parameters arbitrary real number for which the fixed point 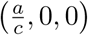 is attracting, repelling and saddle respectively and the three dimensional spaces of *a*, *c* and *e* are plotted in *Fig. 23.*

Here we present examples of attracting and saddle trajectories of the fixed point 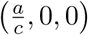.

**Figure 23.**
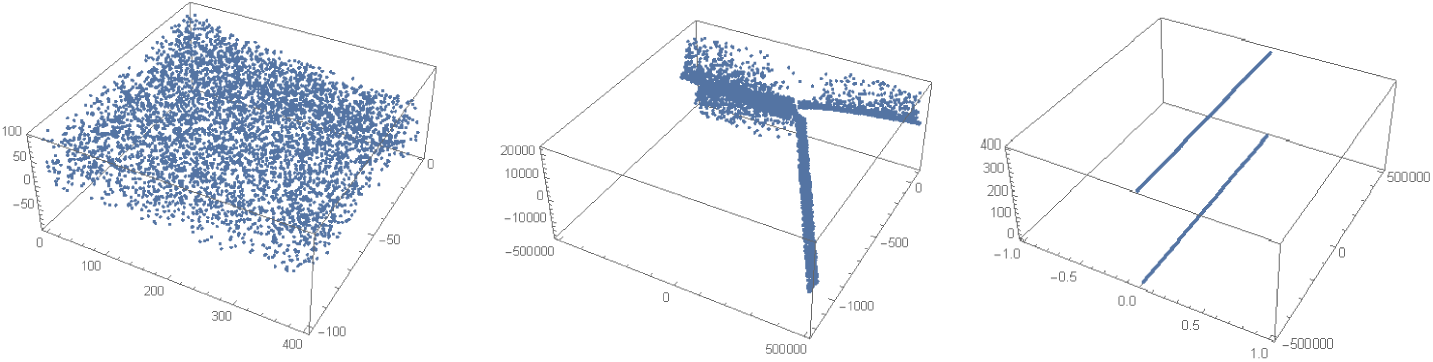
Parameter space of (*a*, *c*, *e*) such that the fixed point 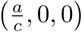 is attracting (left), repelling (middle) and saddle (right) respectively.

First we consider *a* = 50 − 91*i*, *c* = −1 and *e* = 0, the trajectories for ten different initial values taken from the neighbourhood of the fixed point are attracting and that is shown in *Fig. 24.*

**Figure 24.**
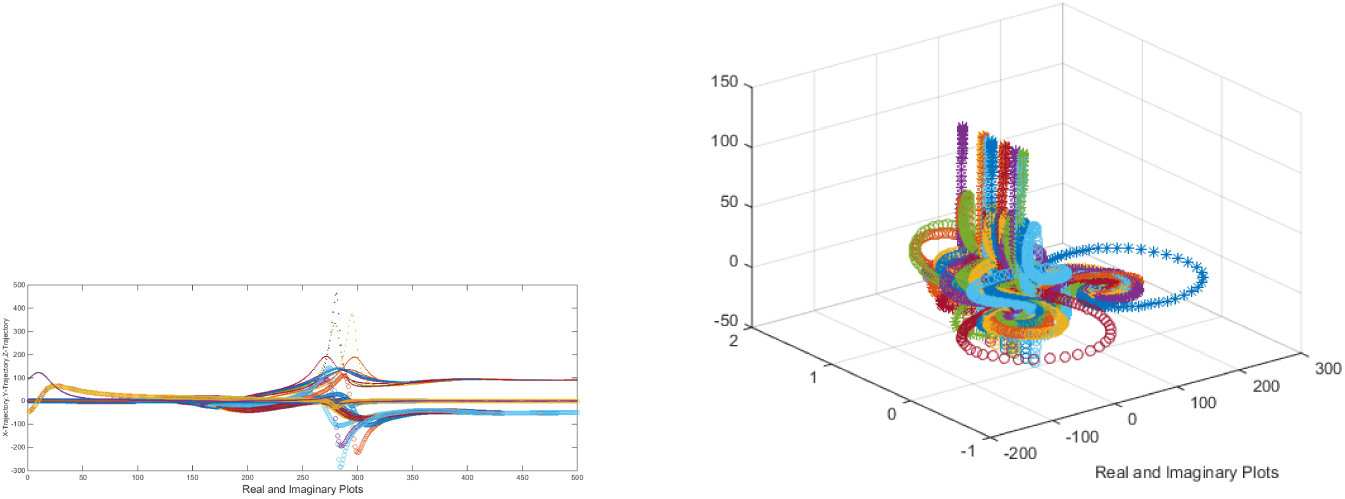
Trajectories attracting towards 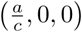

Secondly, we consider *a* = 98.4615 − 172.308*i*, *c* = −8 − 4*i* and *e* = −0.625 + 6.5994*i*, the trajectories for ten different initial values taken from the neighbourhood of the fixed point is saddle and that is figured out in *Fig. 25.*

**Figure 25.**
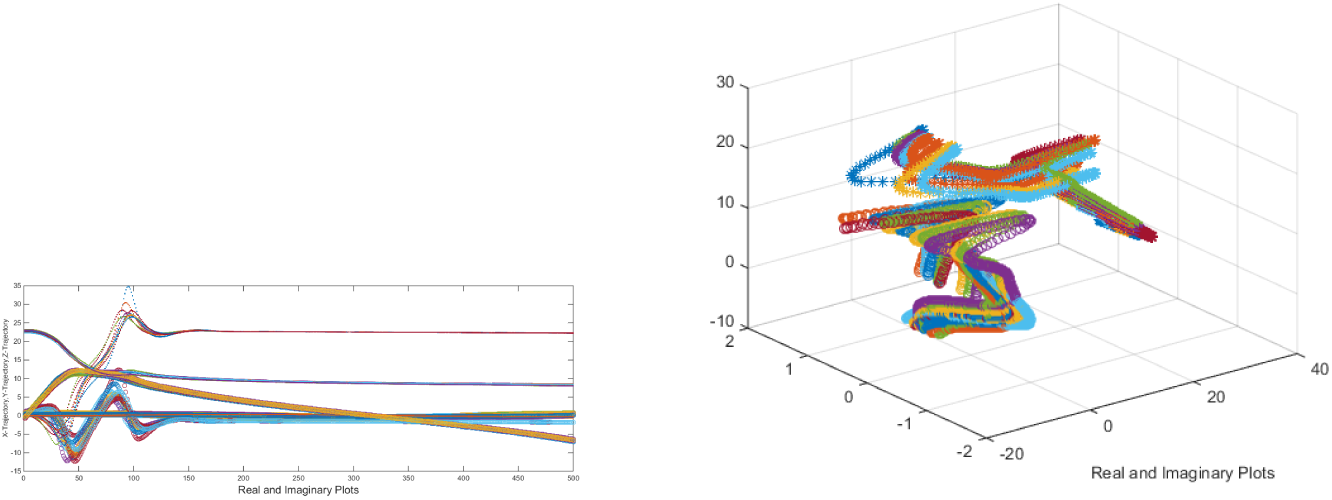
Saddle trajectories attracting of the fixed point 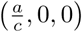

### 3.4. Local Asymptotic Stability of the 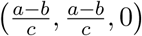

The jacobian about the fixed point 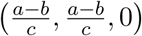 is

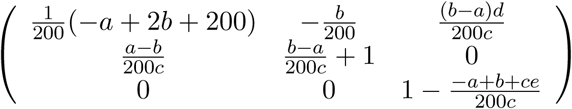

In this consequence, two important theorems are stated below.

#### Theorem 3.8.

*For all the real parameters a, b, c, d and e* (*with dt* = 0.0005), *the fixed point* 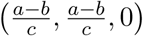 *is attracting*

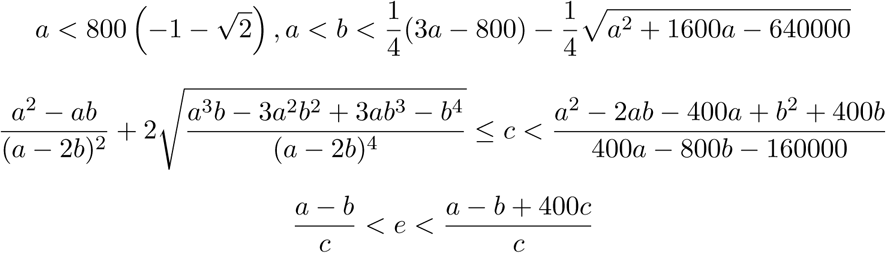

Here we find the two dimensional spaces of parameters (*a*, *b*) and (*c*, *e*) such that the fixed point is 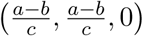 is attracting and the spaces are figured out in *Fig.26.*

**Figure 26.**
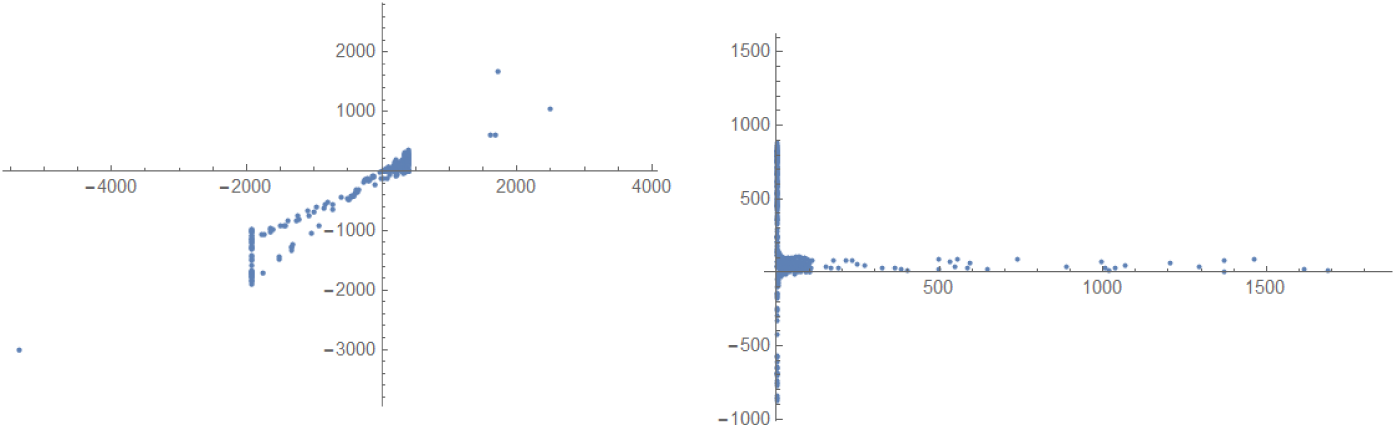
Parameter spaces of (a,b) (left) and (c,e) (right) respectively such that 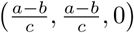 is attracting.

#### Theorem 3.9.

*For all the real parameters a*, *b*, *c*, *d and e (with dt* = 0.0005*), the fixed point* 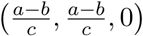 *is repelling*

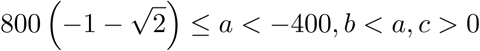

Here we find the three dimensional space of parameters (*a*, *b*, *c*) and (*c*, *e*) such that the fixed point is 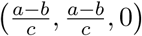 is repelling and the spaces are figured out in *Fig. 27.*

**Figure 27.**
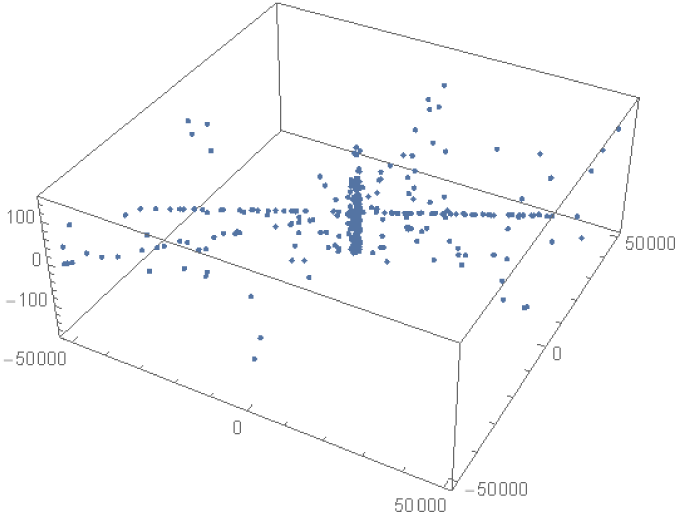
Parameter space of (a,b,c) such that 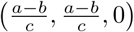 is repelling.

#### Theorem 3.10.

*For all the real parameters a, b, c, d and e (with dt* = 0.0005*), the fixed point* 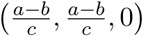 *is a saddle*

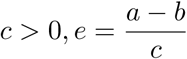

Here we find the three dimensional space of parameters (*a*, *b*, *c*) and (*c*, *e*) such that the fixed point is 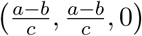 is a saddle and the spaces are figured out in *Fig. 28.*

**Figure 28.**
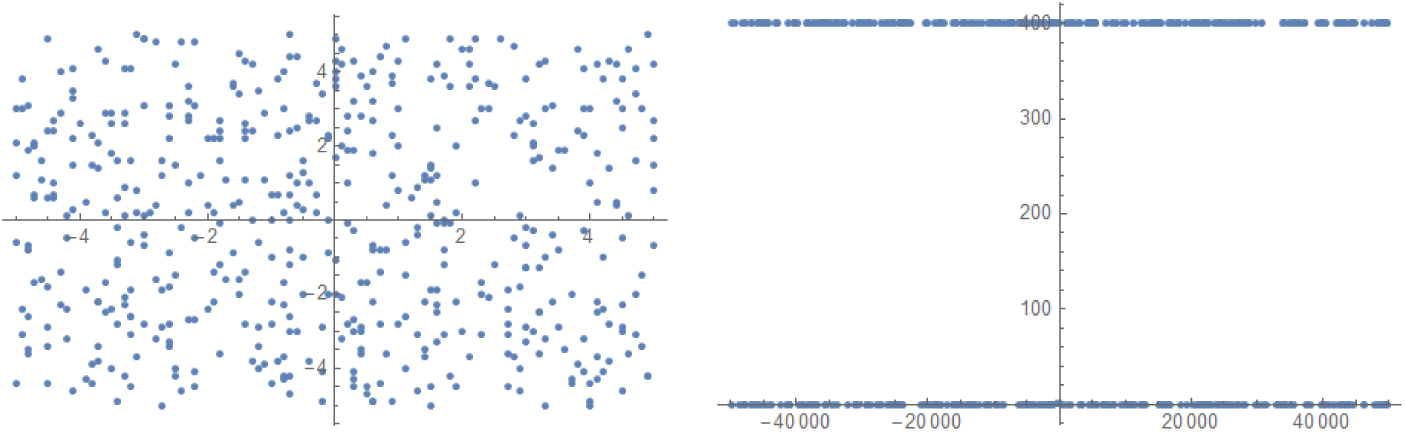
Parameter spaces of (*a*, *b*) (left) and (*c*, *e*) (right) respectively such that 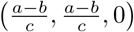 is a saddle.

### 3.5. Local Asymptotic Stability of the 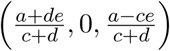

Here we assume the parameter *e* = 0 then the fixed point becomes 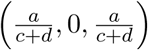 and consequently the density of the eventual population of the immature prey and predator will be same. We wish to see the condition local stability of this fixed point.

#### Theorem 3.11.

*For the real parameters a*, *b*, *c*, *d*, *and e* = 0, *the fixed point* 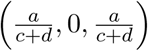 *is attracting if*

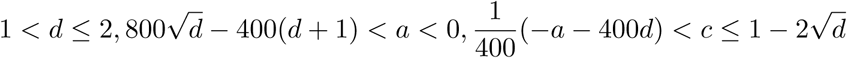

#### Theorem 3.12.

*For the real parameters a, b, c, d, and e* = 0, *the fixed point* 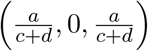 *is repelling if*

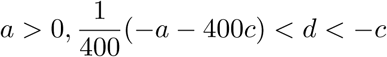

#### Theorem 3.13.

*For the real parameters a, b, c, d, and e* = 0, *the fixed point* 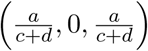 *is a saddle if*

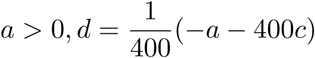

Here we find the three dimensional space of parameters (*a*, *b*, *c*) such that the fixed point is 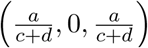 is attracting, repelling and saddle and the spaces are figured out in *Fig. 29.*

**Figure 29.**
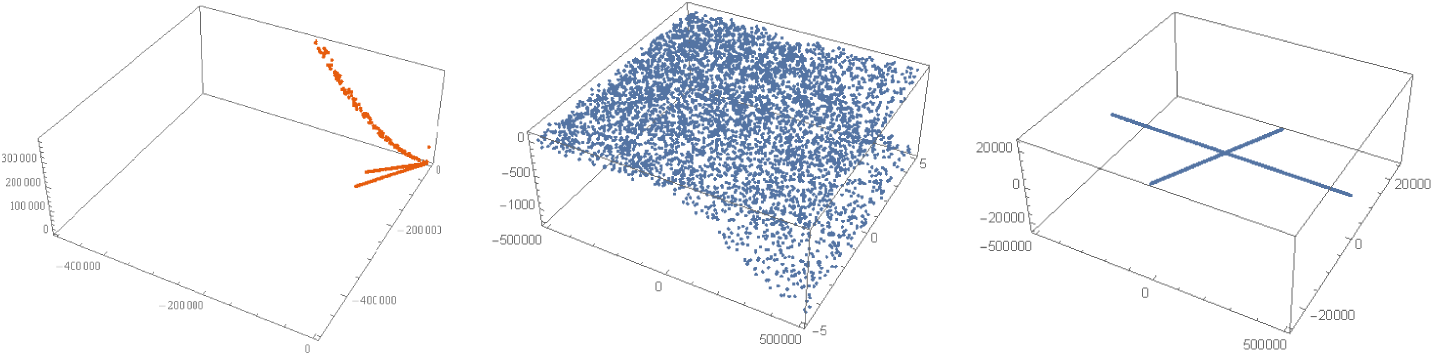
Parameter space of (*a*,*b*,*c*) such that 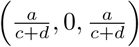 is attracting (left), repelling (middle) and saddle (right) respectively.

Here it is noted that, the fixed point 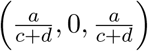 is mostly repelling as we observe from the parameter spaces as shown in *Fig. 29*.

### 3.6. Local Asymptotic Stability of the 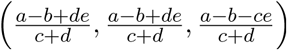

As we did earlier, we fix the parameters and see different cases of fixed points. First we consider *b* = *e* = 0.

#### Theorem 3.14.

*For all real parameters a, c, d with b* = *e* = 0, *the fixed point* 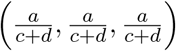 *is attracting if*

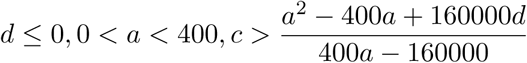

#### Theorem 3.15.

*For all real parameters a, c, d with b* = *e* = 0 (*dt* = 0.0005), *the fixed point* 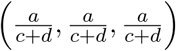 *is repelling if*

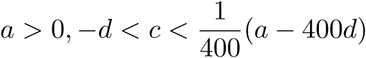

#### Theorem 3.16.

*For all real parameters a*, *c*, *d with b* = *e* = 0 (*dt* = 0.0005), *the fixed point* 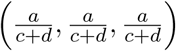 *is a saddle if*

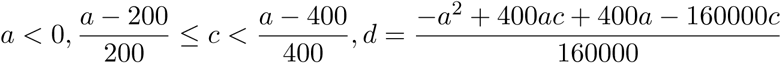

Here we find the three dimensional space of parameters (*a*, *b*, *c*) and (*c*, *e*) such that the fixed point is 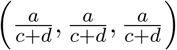 is attracting, repelling and saddle and the spaces are figured out in *Fig. 30.*

**>Figure 30.**
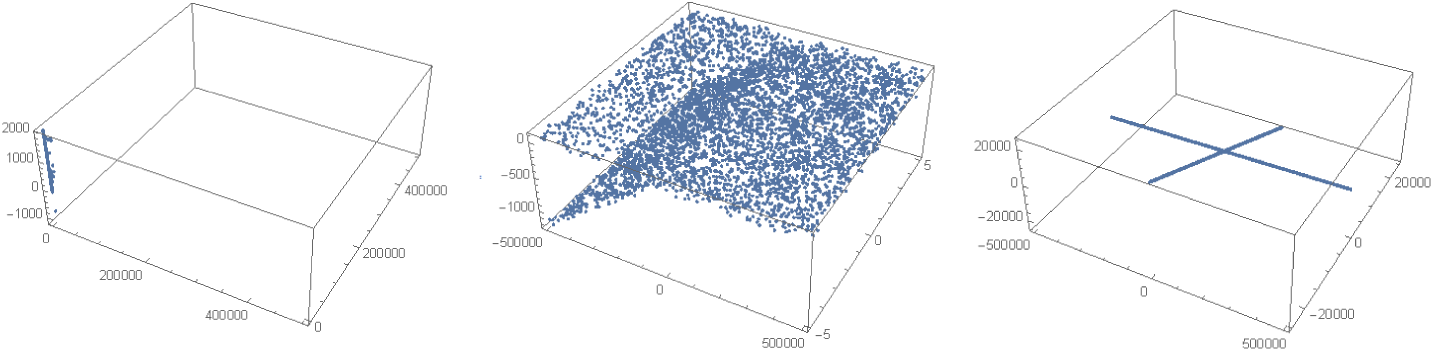
Parameter space of (a,b,c) such that 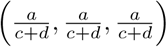 is attracting (left), repelling (middle) and saddle (right) respectively.

Secondly, we consider as we did previously that all the parameters *a*, *b*, *c*, *d* and *e* are equal to *ħ* and then the fixed point becomes 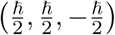.

#### Theorem 3.17.

*There does not exist any real parameters which are all equal such that the fixed point* 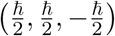 *is attracting.*

*Proof.* The above theorem is seen valid computationally.

#### Theorem 3.18.

*For all parameters a, b, c, d and e equal to ħ while dt* = 0.0005, *the fixed point is* 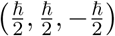 *is repelling if ħ* < −27.823 *or* −2.9806 < *ħ* < 0 *or* 0 < *ħ* < 4.46943 *or ħ* > 28.8255

#### Theorem 3.19.

*For all parameters a*, *bc*, *d and e equal to ħ while dt* = 0.0005, *the fixed point is* 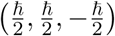 *is a saddle if ħ* = 0 *or ħ* = −27.823 *or ħ* = 28.8255

## 4. Discussions and Future Endeavours

In this paper, the prey (immature and mature) with two stage structures and predator dynamics is reinvestigated with the notion of coupled-dynamics through complex variables. The dynamics of the prey-predator in complex variables is much complicated, than the classical real model, some of the glimpses we observe through computational study. Further a modification is made to the previous model to explore other permanence of prey and predator population. The stability in the modified model are also adumbrated. The present system does not qualify to adhere chaos and higher periodic solutions due to absence of delay in the model. In our future endeavours, we wish to explore the same prey and predator dynamics with delay so that richer dynamics we could experience. This study will help to understand the interpretation of biological phenomena in theory.

## Acknowledgement

The authors tender thank to the Pingla Thana Mahavidyalaya, Maligram, Paschim Midnapur, 721140, India for facilitating scopes in conducting the present research work successfully.

